# Time-lapse mesoscopy of *Candida albicans* and *Staphylococcus aureus* dual-species biofilms reveals a structural role for the hyphae of *C. albicans* in biofilm formation

**DOI:** 10.1101/2023.08.31.555792

**Authors:** Katherine J. Baxter, Fiona A. Sargison, J. Ross Fitzgerald, Gail McConnell, Paul A. Hoskisson

**Affiliations:** Strathclyde Institute of Pharmacy and Biomedical Sciences, University of Strathclyde, 161 Cathedral Street, Glasgow, G4 0RE, UK; The Roslin Institute and Edinburgh Infectious Diseases, Royal (Dick) School of Veterinary

**Keywords:** *Candida albicans*, *Staphylococcus aureus*, Biofilm, Co-Infection

## Abstract

Polymicrobial infection with *Candida albicans* and *Staphylococcus aureus* may result in a concomitant increase in virulence and resistance to antimicrobial drugs. This enhanced pathogenicity phenotype is mediated by numerous factors including metabolic processes and direct interaction of *S. aureus* with *C. albicans* hyphae. The overall structure of biofilms is known to contribute to their recalcitrance to treatment, however the dynamics of direct interaction between species and how it contributes to pathogenicity is poorly understood. To address this, a novel time-lapse mesoscopic optical imaging method was developed to enable the formation of *C. albicans/S. aureus* whole dual-species biofilms to be followed. It was found that yeast-form or hyphal-form *C. albicans* in the biofilm founder-population profoundly affects the structure of the biofilm as it matures. Different sub-populations of *C. albicans* and *S. aureus* arise within each biofilm as a result of the different *C. albicans* morphotypes, resulting in distinct sub-regions. These data reveal that *C. albicans* cell morphology is pivotal in the development of global biofilm architecture and the emergence of colony macrostructures and may temporally influence synergy in infection.

## 1. Introduction

Biofilms are communities of microorganisms within a self-generated extracellular matrix [Penesyan *et al* (2021); Flemming and Wuertz (2019)]. Organisms within biofilms exhibit enhancement of survival against deleterious agents due to the complex matrix of secreted proteins, lipids, extracellular DNA and polysaccharides that surround them [Yin *et al* (2019)]. This complex interaction of organisms and matrix components provides protection against biotic and abiotic stress that is not available to planktonic cells [Bamford, MacPhee and Stanley-Wall (2023); Olsen, (2015)]. In healthcare situations, biofilm-growth can promote survival through limiting diffusion, sequestering, and inactivating antimicrobial agents, along with resistance to mechanical removal [Doroshenko *et al* (2014); Rather *et al* (2021); Peterson *et al* (2015)]. Biofilms may also promote the evolution of antimicrobial resistance through reducing effective concentrations of antimicrobial agents [Allegrucci and Sawer (2008); France et al (2019)] and by facilitating horizontal gene transfer in cells in close contact [Michaelis and Grohmann (2023)].

The dimorphic fungus *Candida albicans* and the bacterium *Staphylococcus aureus* are two prominent opportunistic pathogenic members of the human skin microflora [Haiko *et al* (2019); Janssen *et al* (2021); Budea *et al* (2022)]. Severity of disease caused by *C. albicans* or *S. aureus* can vary from mild cutaneous infection [Di Domenico *et al* (2019)] to systemic infection with multiple tissue involvement [Kwiecinski and Horswill (2020)]. *C. albicans* and *S. aureus* often coinfect [Todd and Peters (2019); Carolus, Van Dyck and Van Dijck (2019)], and become associated with indwelling medical device infections and failure of orthopaedic implants [Qu et *al* (2020); Qu, Peleg and McGiffin (2021); Fernandes and Dias (2013); Cobo et al (2017)]. Several studies have highlighted the propensity of *C. albicans* to enhance *S. aureus* virulence due to activation of toxin production [Todd *et al* (2019), Todd, Noverr and Peters (2019)], increased biofilm growth through prostaglandin production [Krause, Geginat and Tammer (2015)], and enhanced antimicrobial resistance due to *C. albicans* augmenting the biofilm matrix [Kong et al (2016)]. *C. albicans* may also facilitate dissemination of *S. aureus* through adherence to hyphae driving systemic infection [Peters et al (2012)].

Remarkably, biofilm formation by *S. aureus* alone is relatively poor [Cassat, Lee and Smeltzer (2007))]. Yet, in mixed species biofilms cell-cell interactions have been shown to enable biofilm formation, with *C. albicans* adhesins playing a role in local neighbourhood interactions, where *C. albicans* acts as a scaffold for deposition of *S. aureus* cells [Harriott and Noverr (2009)]. The formation of microcolonies of *S. aureus,* coated in secreted matrix derived from *C. albicans,* has been proposed within these mixed species biofilms as the mechanism for increasing antimicrobial resistance in *S. aureus* but not *C. albicans* [Harriott and Noverr (2009)]. Given the role played by *C. albicans* in nucleating *S. aureus* in mixed biofilms and the association of the dimorphic switch with virulence and host-interaction response (Soll (2014), suggests the function played by *C. albicans* cell morphotypes in biofilm structure and dynamics may be vital to understanding mixed biofilm dynamics, yet is currently poorly understood.

To overcome the resolution limitations and inability to resolve macrostructures associated with traditional optical microscopy, the Mesolens can be exploited to image whole mixed species biofilms. The Mesolens bridges the gap in scale between a macrophotography setup and a light microscope [McConnell et al (2016)], with an objective providing the unusual combination of low magnification (4x) and high numerical aperture (0.47). Imaging of specimens up to 6 mm x 6 mm x 3 mm in size can be achieved with a lateral resolution of 700 nm and an axial resolution of 7 µm. Multi-channel imaging is also possible using spectral filtering. Applications of the Mesolens in microbiology have previously revealed the presence of intracolony channels in *E. coli* and their adaptive response to nutrient availability [Rooney *et al* (2020), Bottura *et al* (2022)], but this work has been limited to single species biofilms only.

To determine the role played by *Candida* morphotypes in the structure and dynamics of mixed species biofilms, dual species biofilms containing either yeast-form or hyphal-form *C. albicans* with *S. aureus* were studied using the first instance of time-lapse mesoscopy. This work demonstrates emergence of distinct subpopulations within of each of the species in the mixed species biofilms and that the nature of these is governed by the morphotype of the *Candida* present, leading to unique and remarkable dynamics and features of the mixed *C. albicans/S. aureus* biofilms.

## 2. Materials and Methods

### Generation of a constitutively expressing mCherry *S. aureus* isolate

Chromosomal integration of mCherry into *S. aureus* strain N315 [Kuroda *et al* (2001)] was performed using the methodology outlined in De Jong *et al*, (2017). Briefly, the mCherry harbouring plasmid pRN111 was isolated from *E. coli* strain DC10B and electroporated into electrocompetent *S. aureus* N315. Transformants containing the plasmid were selected on tryptone soy agar (TSA) containing 10 μg/mL chloramphenicol (Cam, Sigma) and incubated for 24-48 h at 30°C. Single crossover events through homologous recombination were generated via two rounds of plating onto TSA containing 7.5 μg/mL Cam followed by incubation at 42°C overnight. Single recombinants were inoculated into 5 ml tryptone soy broth without antibiotics and incubated at 30°C, 200 rpm overnight. Cultures were diluted 1 in 1000 for 5 passages to promote double crossover events. Planktonic bacteria were plated onto TSA with 200 ng/mL anhydrotetracycline (ATc, Sigma) and incubated overnight at 37°C. Integrated mutants were screened by patch plating colonies onto both TSA with and without 10 μg/mL Cam. Colonies that demonstrated the correct phenotype were screened for successful mCherry integration by PCR using primers mCherry_OUT_F: TACGACAATTCAAGAGCTTGC and mCherry_OUT_R: GAGTAAGCCAGAACAGTTCC alongside Whole Genome Sequencing (MicrobesNG, Birmingham UK).

### Strains and Growth Conditions

Fluorescent derivatives of *C. albicans* (pACT1-GFP [Barelle et al (2004)]; constitutive Green Fluorescent Protein [GFP] expressing) and *S. aureus* (N315 constitutive mCherry expressing,(described above) were cultured on YPD and LB agar respectively.

Seed cultures for dual species biofilm were generated from single colonies of each species inoculated as monocultures in 5 ml of Lysogeny Broth (Merck Life Science, UK) supplemented with 0.2% glucose. Those 5 ml overnight cultures destined for hyphal-form biofilms were incubated for 16 hours with shaking at 37 °C /250rpm, resulting in hyphal formation within the *C. albicans* stationary phase culture.

Overnight cultures destined for yeast-form biofilms underwent a dilution step in culture set-up to allow *C. albicans* to reach stationary phase prior to hyphal formation. Seed cultures for yeast-form biofilms were initially inoculated identically to those for hyphal-form biofilms, however, directly after inoculation, those 5ml monocultures were vortexed to disperse cells, and immediately used to inoculate a second set of 5 ml Lysogeny Broth at a dilution of 500-fold for *C. albicans* monocultures, and1000-fold for *S. aureus* monocultures. These diluted monocultures were used as yeast-form seed cultures, and were incubated for 16 hours with shaking at 37 °C /250rpm. After the 16 hours, viable cells (colony forming units; CFUs) were determined from each culture by plating prior to the culture mixing step (described below)

### Time lapse mesoscopy of biofilm growth

Biofilm growth conditions were adapted from the colony spreading biofilm methodology described by Kaito and Sekimizu (2007). The 16 h cultures (see above) were mixed and 2 µl spotted on to LB agar mesoscopy mounts (see below), air-dried for 10 minutes, and immediately imaged to produce time zero images of initial cell deposition. Initial spots consist of averages of 3.83 x10^9^ CFU/ml *S. aureus* and 1.3 x10^8^ CFU/ml *C. albicans.* After initial t=0 imaging, samples were incubated at 37 °C for 12 hours with removal at 3, 6, 9 and 12 hours post-inoculation for time-lapse imaging of biofilm formation. After imaging at the indicated time point, samples were returned to the incubator.

### Development of a specimen mount for time lapse mesoscopy

Imaging of samples for optical mesocopy is best achieved in optically matched liquid (McConnell *et al* (2016)), however colonies of both *C. albicans* and *S. aureus* dissipate when immersed in liquid, negating the use of previously established Mesolens imaging methods (Rooney *et al* (2020)). Imaging through a thin layer of LB agar solidified on a glass coverslip permitted biofilms to be viewed through the base of the sample (Figure 1A & B), facilitating water immersion for best image quality, and allowing biofilm formation to be followed over time. (Supplemental Figure 1-Image of mount). Samples were prepared and followed over time as described above.

**Figure 1:**
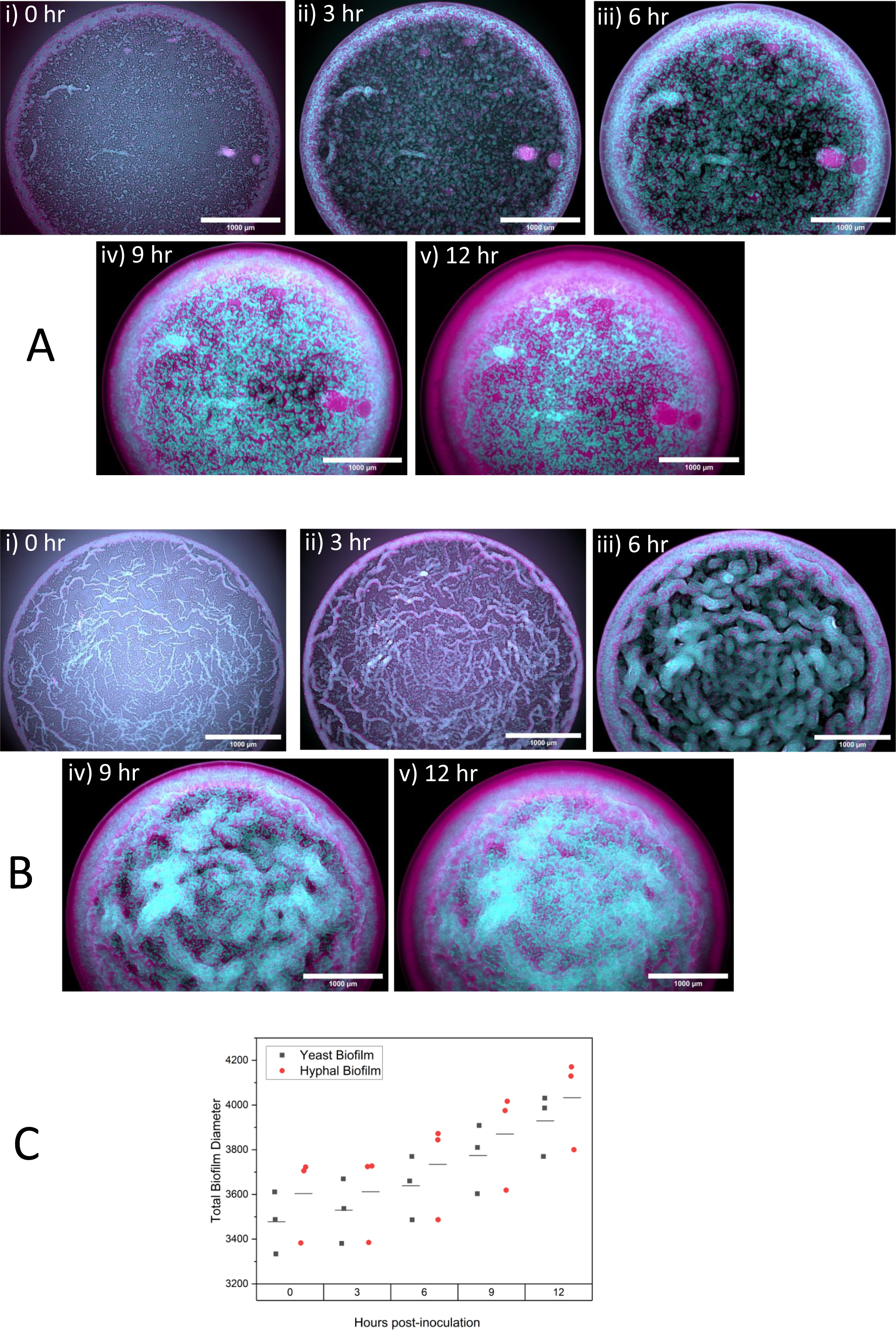
Time-lapse optical mesoscopic imaging. 12 hour time-lapse Mesoscopy of *C. albicans* pACT-1 GFP (cyan) */S. aureus* N315 mCherry (magenta) dual species biofilms. Panel A: Yeast-form *C. albicans/S. aureus.* Panel B: Hyphal-form *C. albicans/S. aureus.* (i) Initial deposition at t=0; (ii) t3 – 3 hours post-inoculation; (iii) t6-6 hours post-inoculation; (iv) t9 – 9 hours post-inoculation; (v) t12= 12 hours post inoculation. Images are maximum intensity z-projections. Scale bar 1000 µm. Graph C: Average diameter of both Yeast-form and Hyphal-form biofilms at each time point.

### Sugru dam manifold manufacture and preparation

Sugru mouldable glue (Sugru, Amazon) was moulded into a long cylinder and adhered to a glass coverslip 70 mm x 70 mm type 1.5 0107999098 (Marienfeld, Lauda-Koenigshofen, Germany) to form a circular dam 45 mm in diameter, and approximately 10 mm in height. After adhesion to the glass coverslip, the circular Sugru dam was left to cure for 24 hours prior to use as per manufacturer instructions. Manifolds were sterilised by immersion in 70% (v/v) ethanol, then air-dried and exposed to UV light for 15 minutes. The interior of the dam was filled with molten LB agar supplemented with 0.2% glucose to a depth of 2 mm, and left to solidify. Manifolds were prepared on the day of imaging, prior to culture mixing.

### Time Lapse Mesoscopy

Widefield epi-fluorescent mesoscopic imaging was performed by modification of the method described by Bottura *et al* (2022). Excitation of fluorescent proteins was achieved with a CoolLED pE-4000 LED (CoolLED, UK) at excitation/emission wavelengths of 490/525 ± 20 nm for GFP and 585/635 ± 20 nm for mCherry. High-resolution images were captured using a chip-shifting camera sensor (VNP-29MC; Vieworks, Anyang, Republic of Korea) which recorded images by shifting a 29-megapixel CCD chip in a 3 x 3 array [Schniete et al]. In this mode, the sampling rate was 4.46 pixels/µm: this corresponded to a 224 nm pixel size, satisfying Nyquist sampling criteria.

Mesoscopy was performed with the correction collars of the Mesolens set for water immersion to match the refractive index of LB agar (1.33 vs 1.34, respectively). To capture the three-dimensional spatial distribution of *C. albicans* and *S. aureus* as biofilms developed over time, z-stacks of biofilms were acquired over a 12 hour time period. To obtain z-stacks, the specimen was moved in the axial direction in 5 µm increments using a computer-controlled specimen stage (Optiscan III, Prior Scientific). Biofilms were incubated at 37°C with imaging performed at time 0 h and at 3 h intervals thereafter.

### Image analysis

Z-stacks were converted to maximum intensity projections using the FIJI image processing software (ImageJ, version 2.1.0/I.53c) [Schindelin *et al* (2012)]. Measurements of *Candida* hyphal width, biofilm core diameter, halo width and total diameter were taken with the measurement feature of FIJI. To understand the degree of association of *S. aureus* with *C. albicans* in biofilm structures, colocalization of the strains was undertaken. Selection of the core region of the biofilm from the halo regions was performed by merging GFP and mCherry channels into a two-colour image, cropping the total available core area within the halo and splitting the cropped image back into separate channels. Each resulting colour channel was subjected to image thresholding, converted to a binary image and subsequently analysed with the ‘Just another Colocalisation’ Plugin (JACoP) [Bolte and Cordelieres (2006)] to determine colocalization of *C. albicans* and *S. aureus* in biofilms. Pearsons’s correlation coefficient (PCC) and Manders’ overlap coefficients (MOC) were used to statistically assess the degree of colocalization. PCC analysis of core regions used the intensities of each signal (mCherry or GFP) for a given pixel across the appropriate colour channel to calculate the correlation of signal intensity of each channel with the other. A PCC value of zero indicates there is no association between the two signals. Increasing positive values indicate increasing levels of colocalization. MOC is a measure of the fraction of overlapping signals, the M1 coefficient is the degree GFP signal overlaps with mCherry signal, and the M2 coefficient is the degree mCherry signal overlaps with GFP. Dual channel images were generated using QiTissue software (Version 1.2.1 Pre-release Version, Quantitative Imaging Systems LLC). To identify whether hyphae were components of the *C. albicans* projections into the *S. aureus* peripheral halo, plot profiles of projections were derived from the FIJI line measurement tool and the resulting grey value curves analysed using the full width half maximum (FWHM) principle, where the width of the curve is measured between the y-axis points that are half the height of the curve. Values were calculated using the FWHM.ijm macro available on GitHub. (https://gist.github.com/lacan/45f865b5a38d7c3a96a7cd8b25923407)

### Antibiotic susceptibility testing of biofilms

Decreasing concentrations of vancomycin (Fisher Scientific, UK) were generated by the broth dilution method (EUCAST (2003)) to establish the lowest concentration of vancomycin required to prevent growth of planktonic *S. aureus* in overnight cultures (3.125 µg/mL). Once identified, a variation of the protocol undertaken by Adam *et al* (2002) was used to test the susceptibility of planktonic seed cultures of *S. aureus* grown in the presence of yeast-form and hyphal-form *Candida* cells after 5 hours exposure to vancomycin. Planktonic cultures (16 hrs) were treated with vancomycin (3.125 µg/mL) and incubated at 37°C with shaking for 5 hours. In the case of dual species biofilms, 6 hour biofilms were treated with 4µl of LB containing 3.125 µg/mL onto the top of each biofilm and allowed to dry before being incubated at 37°C for 5 hours. After exposure, biofilms were removed from agar by scalpel and placed into an Eppendorf tube, resuspended v/v into 1 mL of LB by mixing. The resulting cell suspension was used to determine viability by CFU count. All sampling was performed in triplicate.

## 3. Results

### Common features of biofilms of both *Candida* morphotypes are the result of fluid dynamics

Initial seeding of the biofilm cultures resulted in founder populations of *C. albicans* and *S.* aureus in yeast-form biofilms averaging 3.65 x10^9^ CFU/ml *S. aureus* and 1.34 x10^8^ CFU/ml *C. albicans,* and 4 x10^9^ CFU/ml *S. aureus* and 1.28 x10^8^ CFU/ml *C. albicans* in hyphal-form biofilms. Evaporation of the mixed cell inoculation mixture resulted in cells being deposited in the archetypal ‘coffee ring’ structure due to fluid dynamics associated with evaporation of liquid drops [Agrawal *et al* (2020)]. This causes the formation of a ring of *C. albicans* and *S. aureus* cells (halo), surrounding a ‘core’ of sparsely deposited cells. This was found in cultures established with both yeast-form and hyphal-form *Candida* in the cell mixture (Figures 1A & B).

### Macrostructures emerge in mixed species biofilms containing either *Candida* **morphotype**

Mesolens images captured over the 12 hour time period reveals the emergence of similar segregation of the species within the halo of both biofilms, and distinct morphotype-dependent core structures. Close inspection of cell morphology at t=0 indicates the presence of only yeast cell morphotype *C. albicans* and *S. aureus* in both Hyphal and Yeast-form biofilm halos [Figure 2A], whereas the presence of hyphae is only observed in the core of the Hyphal-form biofilm [Figure 1A (i) vs Figure 1B (i)].

**Figure 2:**
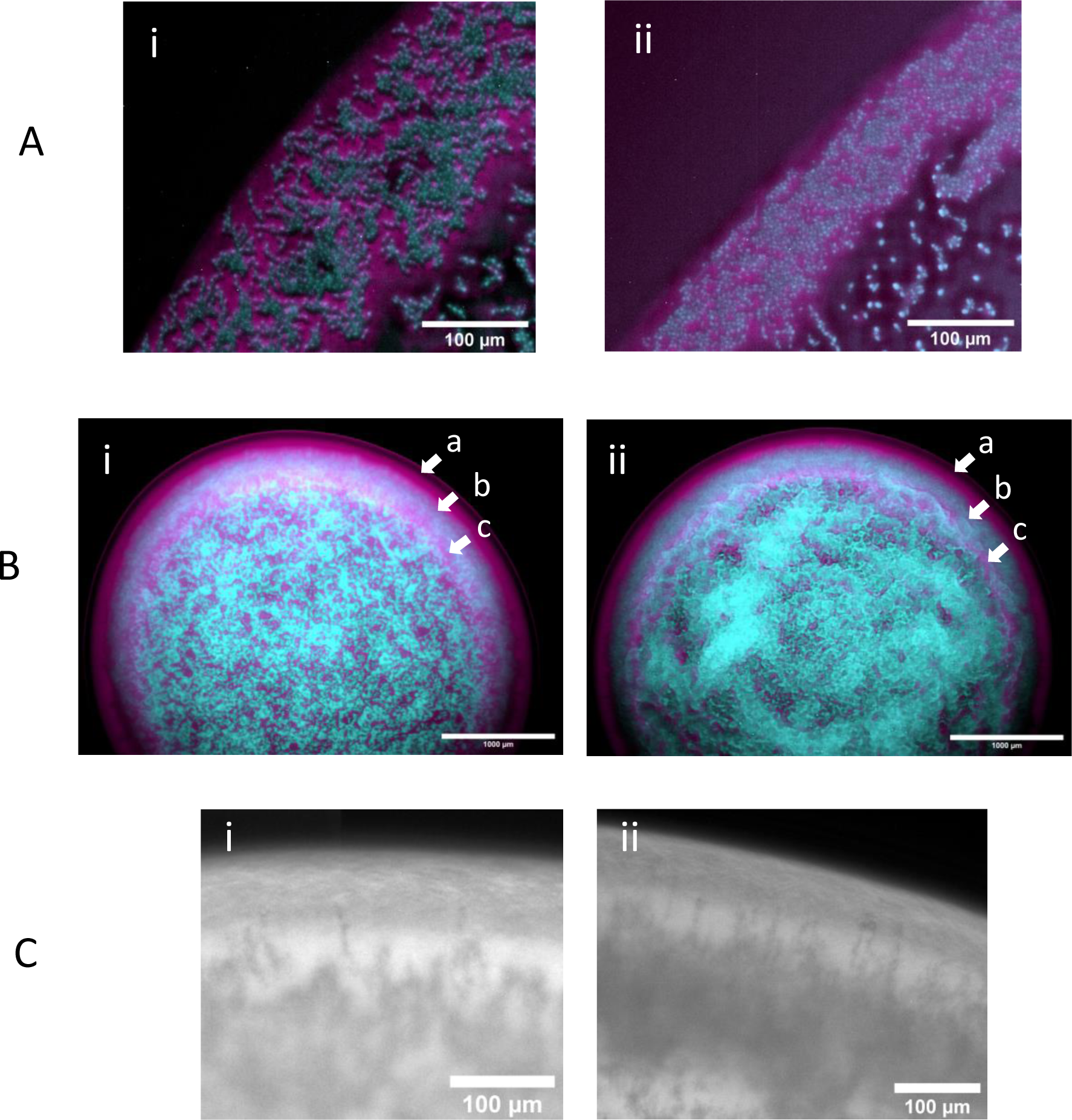
Yeast-form and Hyphal-form halo structures of pACT-1 GFP (cyan) */S. aureus* N315 mCherry (magenta) dual species biofilms. **Panel A**: Digital zoom of halo illustrating similar cellular form and composition at initial deposition (t=0h) in (i) Yeast-form *C. albicans/S. aureus* biofilm and (ii) Hyphal-form biofilm. **Panel B**:Comparison of species banding patterns in halos of (i) Yeast-form *C. albicans/S. aureus* biofilm and (ii) Hyphal-form biofilm 12 hours post-inoculation (t=12h). All images are (or are derived from) maximum intensity z-projections. Scale bar in A: 100 µm; scale bar in B: 10000 µm; scale bar in C: 1000 µm. **Panel C:** Digital zoom of halo illustrating serrated interface between *C. albicans* and peripheral *S. aureus* halo bands as hyphae penetrate *S. aureus* band in (i) Yeast-form *C. albicans/S. aureus* biofilm and (ii) Hyphal-form biofilm 9 hours post-inoculation (t=9h) Scale bar in C: 100 µm.

In the early stages of biofilm development, *C. albicans* and *S. aureus* within the halos of both *C. albicans* morphotype biofilms begin to emerge into discrete species-specific banding patterns. These develop into a middle band of *C. albicans* flanked by an outer band of *S. aureus* on the periphery of the biofilm, and a broken discontinuous internal band of *S. aureus* on the core-proximal edge of the halo by 12 hours [Figure 2B]. This structure is first noticeable as an internal band of *C. albicans* cells at the 3 hour time point, followed by the emergence of the faster growing *S. aureus* as a band at the periphery by 6 hours, as shown in Figure 1A (ii-iii), and 1B (ii-iii) respectively. In later stages of biofilm formation at 9 and 12 hours post-inoculation, an additional broken band of *S. aureus* emerges at the core-facing edge of the halo [Figures 1A (iv-v) and 1B (iv-v))]. At 9 hours, a serrated boundary is observed between the internal *C. albicans* band and the peripheral *S. aureus* band, with multiple fingers of *C. albicans* invading the peripheral subpopulation of *S. aureus* [Figure 2C]. These *C. albicans* incursions begin as hyphae from the *C. albicans* halo band, as their starting width at t=6 hours is between 2-4 µm. [Supplemental figure 2]. By 12 hours, similar halo banding patterns are observed in biofilms of both *C. albicans* cell morphotypes [Figure 1A (v) and Figure 1B (v)].

Strikingly, core structure is *C. albicans* cell morphotype dependent. There is a prominent difference in cell deposition between the Yeast-form and Hyphal-form dual species biofilm cores, noticeable from the initial time point. In the Yeast-form biofilm, cells of both *C. albicans* and *S. aureus* are randomly deposited within the core at deposition [Figure 3 (i)] However, cells from the Hyphal-form culture accumulate in aggregates of *S. aureus* and *C. albicans* yeast-form cells clustered around hyphae, with larger cell-free gaps between these cell groupings. This differential deposition results in the development of two architecturally distinct core structures as the biofilms mature. At 3 hours post inoculation in the Yeast-form biofilm, *C. albicans* and *S. aureus* cells begin to develop into discrete microcolonies of each species [Figure 1a(ii)], which cover the core with a mosaic-like pattern by the 12 hour time point [Figure 1a (v)]. This is in marked difference to the Hyphal-form biofilm, where the hyphal-cell aggregates expand into ropes of intercalated *C. albicans* and *S. aureus* over the first 6 hours post inoculation [Figure 1b (ii)], resulting in a marbled-like distribution of *C. albicans* and *S. aureus* by t=12 [Figure 1b (v)].

**Figure 3:**
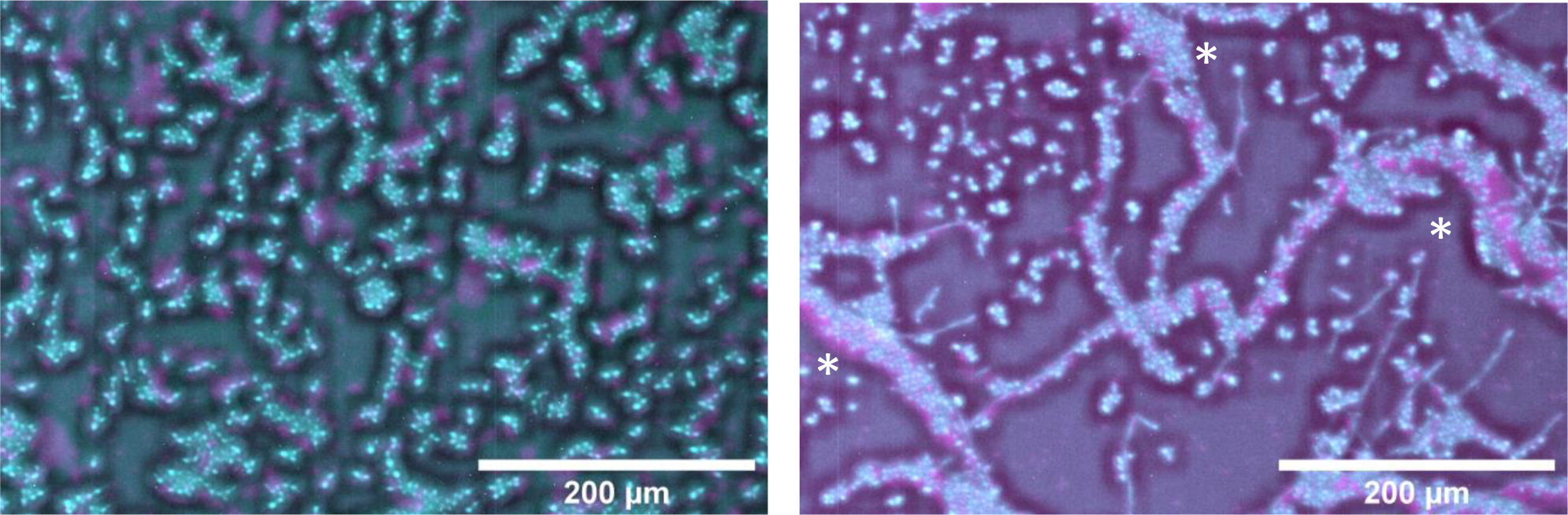
Impact of *C. albicans* cell morphotype on cell deposition within biofilm cores of pACT-1 GFP (cyan) */S. aureus* N315 mCherry (magenta) dual species biofilms. (i) Digital zoom of Yeast-form biofilm and (ii) Hyphal-form biofilm at t=0. Asterisks in (ii) indicate *S. aureus* and Yeast-form *C. albicans* accumulation along hyphal-cell aggregates

### Quantification of cell-derived macrostructures reveals impact of *C. albicans* cell morphotype on biofilm architecture

Following initial deposition, there are noticeable differences in the cell-derived macrostructures of the hyphal-form biofilm and the yeast-form biofilm in terms of halo width, core diameter, and deposition of cells within the core. We quantified these parameters to understand the dynamics of these macrostructures and their development [Figure 4]. Both types of biofilms have similar total biofilm diameters at all time points [Figure 4B], averaging 3930 µm (± 106) for yeast-form biofilms and 4033 µm (± 155) for hyphal-form biofilms at the 12 hour time point. Biofilm expansion rates are comparable for each biofilm type, averaging between 3.6-3.9 µm/hr between 3-12 hours post-inoculation [Supplementary Figure 3]. The halos of hyphal-form biofilms are noticeably smaller than those of yeast-form biofilms, with an average a width of 100 µm (± 13) compared to 180 µm (± 7) respectively at deposition. Differences in halo width between the hyphal-from and yeast-form biofilms decreases over the growth period, however hyphal-form biofilm halos remain narrower than those of yeast-form biofilms, with an average of 390 µm in comparison to the larger yeast-form halo average of 430 µm at 12 hours.

**Figure 4:**
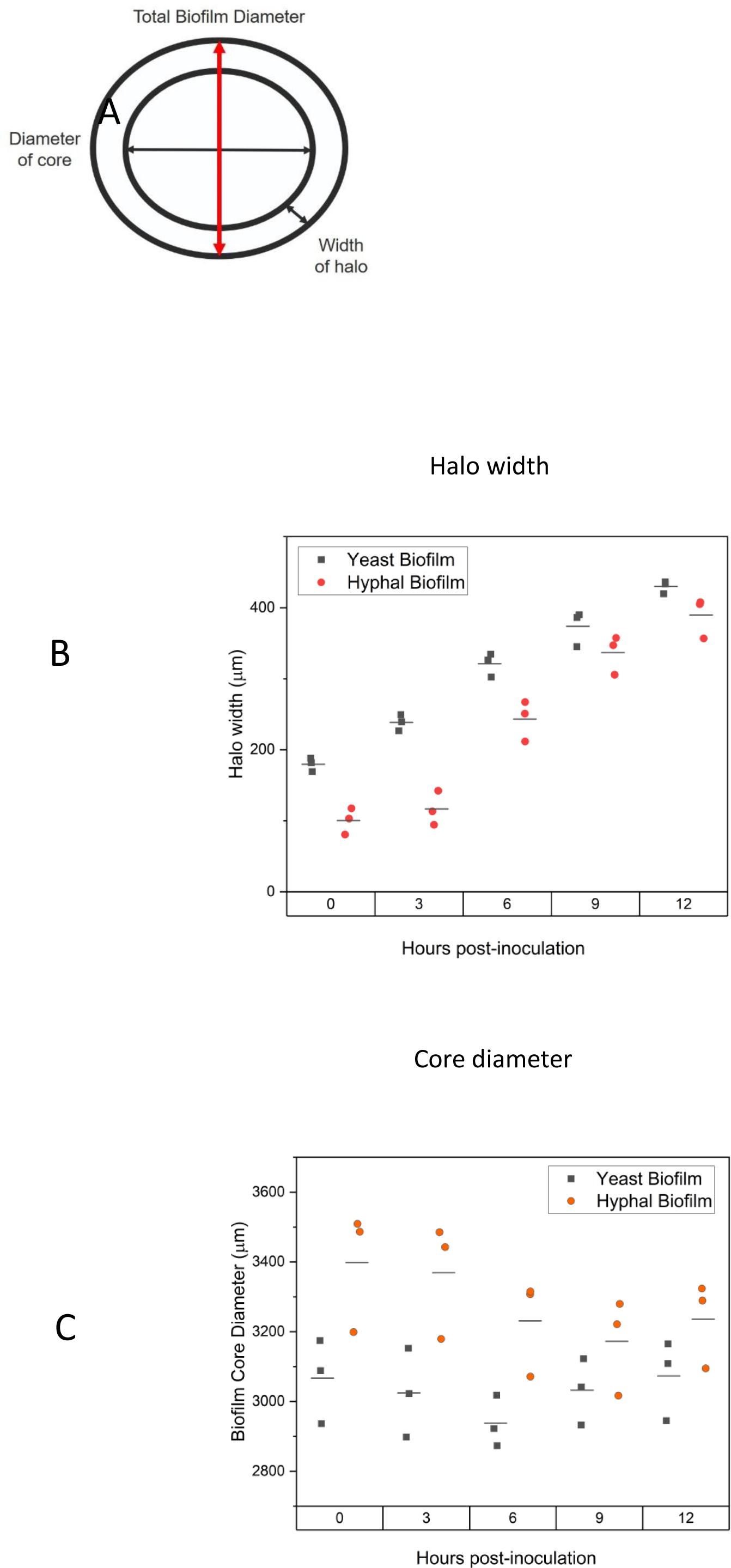
Quantification of Yeast-form and Hyphal form biofilm macrostructre properties from time-lapse mesoscopy images. For each time point, three measurements of each property were obtained from three independent biofilms (n=9). Standard deviation error bars. **Panel A:** Diagram of properties measured **Graph B:** Halo width of Yeast-from (green diamonds) or Hyphal form (orange crosses) biofilms. **Graph C:** Core diameter of Yeast-from (green diamonds) or Hyphal form (orange crosses) biofilms. P values (supplemental data) indicate these results are significant.

As both biofilms have similar total diameters at all time points, we postulated that our findings of a smaller halo width would also imply a larger core diameter as these two components together compose the total biofilm diameter. Using the same FIJI image analysis method, the quantification of core diameter indicates that this is the case (Figure 3). The core of hyphal-form biofilms at deposition average 3398 µm (± 133) versus the smaller Yeast-form diameter at 3067 µm (± 87). The Hyphal-from consistently has a larger diameter than that of the Yeast-form biofilm across the 12 hours. Again, as for the halo data, the difference between core diameters of hyphal-form and yeast-form biofilms does decrease over time, however this difference is still significant (statistical analysis in supplementary table 1) and may provide an advantage in infection, as a larger core will provide a greater area for colonisation, and increase gas exchange and nutrient access for cells in a Hyphal-form biofilm than in a Yeast-form biofilm.

### Aggregation of free cells by *C. albicans* cells leads to greater interspecies clustering in hyphal-form biofilms

Due to the correlation of hyphal invasion with systemic *S. aureus* infection [Schlecht *et al* (2016)], we undertook analysis of co-localisation of mCherry and GFP signals within biofilm cores to understand the degree of association of *S. aureus* with yeast-form and hyphal-form *C. albicans* in global biofilm structure. In our imaging, we can see association of *S. aureus* with hyphae at time deposition in line with previous findings [Harriett and Noverr (2009); Peters *et al* (2010); Peters *et al* (2012); Kean *et al* (2017); Zago *et al* (2015)]. It was also noted that *S. aureus* accumulate along the edges of hyphal-cell aggregates (asterisks Figure 3i). This suggests there is more to the *S. aureus/C. albicans* physical relationship than *S. aureus*-hyphal interaction alone. It was hypothesised that hyphae may function as a nucleation point for free yeast-form *C. albicans* and *S. aureus* in biofilm. To quantify this observation of increased association of *C. albicans* and *S. aureus* in the core of the hyphal-form than in the yeast-form biofilm, Pearson’s Coefficient and Manders Coefficient were used to quantify correlation and co-occurrence of both species using the JACop FIJI plugin [Bolte S and Cordelieres] [Figure 5(a)]. It was apparent that Hyphal-form biofilms have both a higher Pearson’s Coefficient [Figure 5(b)] and higher Manders coefficients [Figures 5(c) and (d)] from the deposition point onwards, indicating *C. albicans* and *S. aureus* localise at a greater incidence and for a longer period of time in the biofilm core when hyphal-form *C. albicans* is present. This correlation occurs until 9 hours of growth (Table 2, supplementary data). However, this is maybe influenced by population densities at t=12, where both cores are densely packed with both *C. albicans* and *S. aureus* structures, and fluorescence may be too great to resolve the relationship.

**Figure 5:**
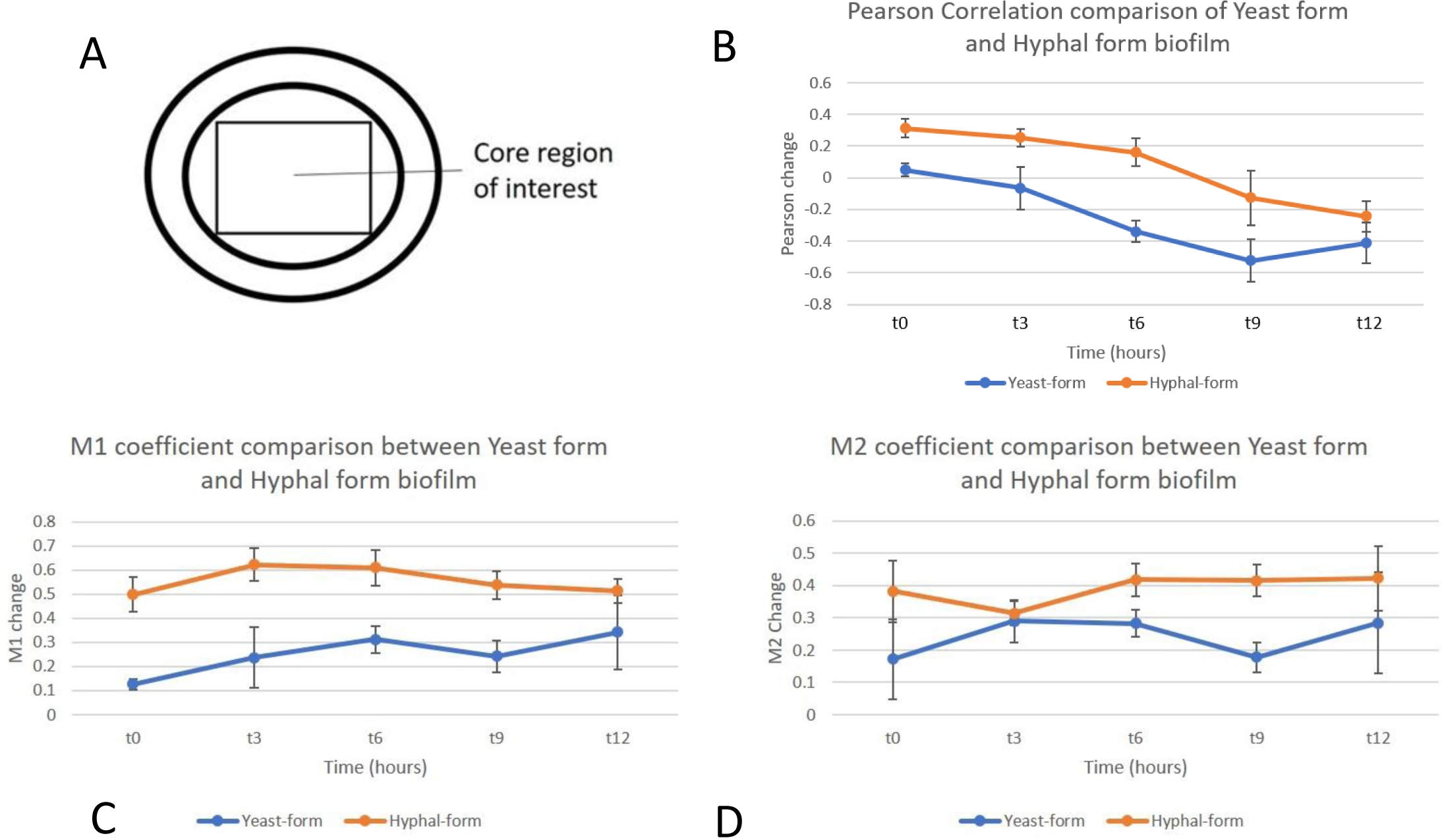
Colocalisation of *C. albicans* and *S. aureus* in Yeast-form and Hyphal-form dual species biofilms from time lapse mesoscopy images. **Panel A:** Diagram indicating core region of interest selected for colocalization analysis. **Graph B:** Comparison of change in Pearson coefficient between Yeast-form (blue) and Hyphal-form (orange) dual species biofilms at each time point, representing the degree of colocalization of *C. albicans* GFP and *S. aureus* mCherry signal. Positive coefficients denote localisation, Negative coefficients denote inverse localisation (i.e, greater signal of one correlating with lower signal of other). **Graps C and D**: Manders coefficients M1 (overlap of GFP signal with mCherry signal) and M2 (overlap of mCherry signal with GFP signal) respectively, representing the degree of overlap of fluorescent signals. The greater the coefficient value, the more overlap between signals at the same locality. Measurements taken from 3 independent biofilm cores of each biofilm subtype, error bars are standard deviation at each time point. P values (supplemental data) indicate these results are significant.

### Biofilms of both *C. albicans* cell morphotypes confer increased *S. aureus* resistance to vancomycin

It is well documented that association with *C. albicans* increases the resistance of *S. aureus* to antibiotics [Carolus, Van Dyck and Van Dijck (2019); Harriot and Noverr (2009); Hu *et al* (2021), Kong *et al* (2016)]. To test if the morphotype of *Candida* present in the biofilms influences antibiotic resistance, vancomycin sensitivity of both *S. aureus* planktonic culture and *S. aureus* biofilm-derived cells from either yeast and hyphal dual species biofilms was tested. Biofilms were tested after 6 hours of growth as this is the point where the macrostructures are established.

After exposure, the cell viability of planktonic *S. aureus* is reduced significantly by a factor of 3000-fold, in comparison, vancomycin treated yeast-form and hyphal-form biofilms do not show a reduction in viability, with similar numbers of viable cells with or without vancomycin treatment [Figure 6]. These findings confirm that even after only 6 hours of biofilm growth facilitates *S. aureus* resistance to vancomycin when compared to planktonic cultures. No difference was observed any difference between the viability of *S. aureus* in yeast-form and hyphal-form biofilm after vancomycin exposure under these conditions.

**Figure 6:**
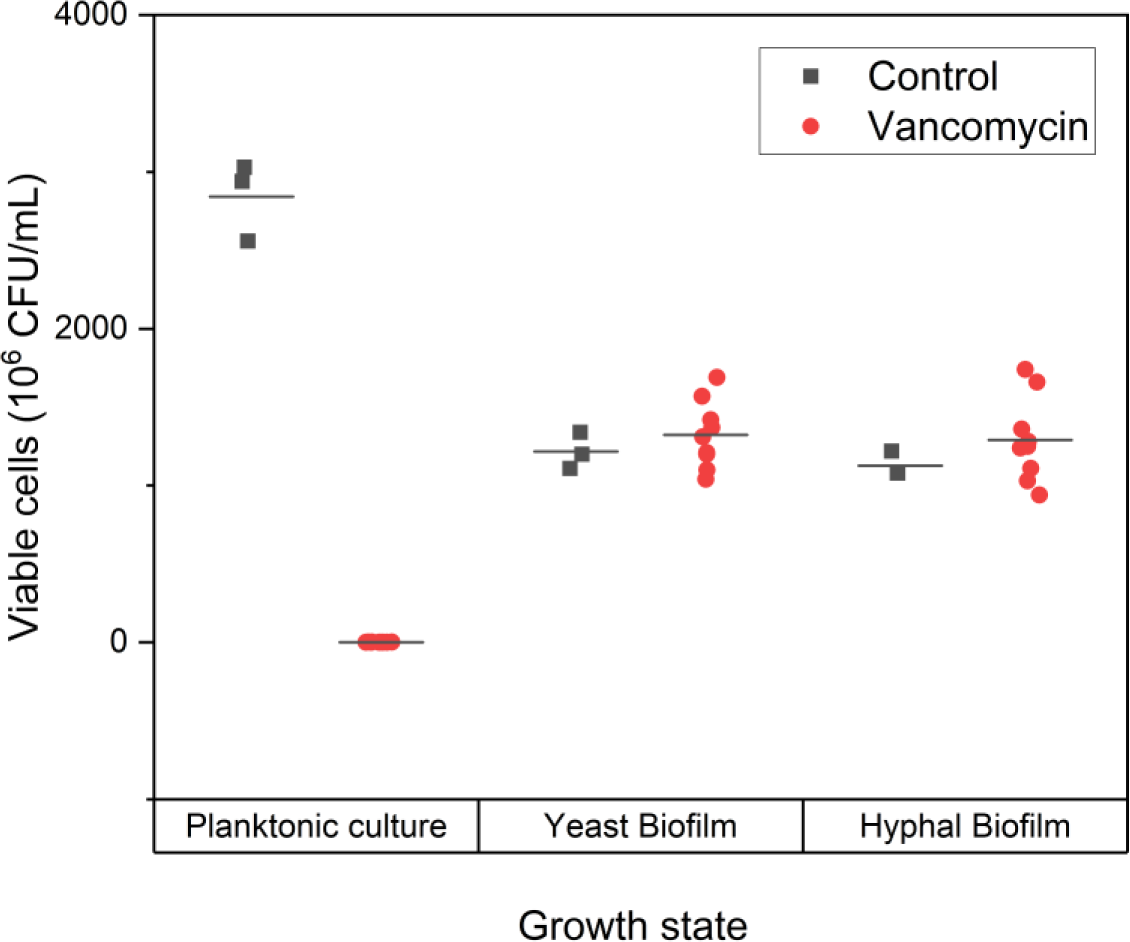
Viability of *S. aureus* after 5 hours exposure to vancomycin. Graph displays viable cells determined by CFU count. Planktonic cells used were *S. aureus* biofilm seed cultures. Data presented as 10^6^ CFU/ml. Biofilms of both *C. albicans* morphotypes confer increased resistance to vancomycin in comparison to planktonic culture.

## Discussion

Dual species biofilms are of great importance in the clinic [Eichelberger *et al* (2023); Bouza *et al* (2013)]. To determine the role played by *Candida* morphotype in the development and dynamics of dual species biofilms a time-lapse mesoscopy method was developed. This enabled the formation of dual species biofilms produced by *C. albicans* and *S. aureus* to be followed in intricate spatial detail, from deposition of individual cells to mature biofilm structure. This provides an unprecedented level of detail on the emergence of cell-governed macrostrucures within the *C. albicans/S. aureus* dual species biofilm.

These data suggest the formation of multiple large interspecies clusters by hyphae facilitates interaction of *C. albicans* and *S. aureus* in biofilms from initial deposition onwards, promoting their physical and metabolic interplay within the core. In these clusters, *C. albicans* matrix components likely coat adjacent *S. aureus* cells from an earlier stage of biofilm formation than in non-hyphal biofilm. These interactions potentially increase the protective effect of β-1,3-glucans from *Candida* against antimicrobial challenge over the lifetime of the biofilm [Kong *et al* (2017)]. Clustering of cells will also increase *S. aureus* exposure to metabolites secreted by *C. albicans*, influencing *S. aureus* expression of the *agr* sytem through *C. albicans* alkalisation of the local environment [Todd *et al* (2019); Todd, Noverr and Peters (2019)], and promoting growth via Prostaglandin E2 [Krause, Geginat and Tammer (2015)]. These effects potentially produce a larger population of *S. aureus* with *C. albicans*-augmented virulence and growth capacity than in biofilms where hyphae are absent, resulting in less associated *C. albicans* and *S. aureus* cells. This physical interaction between the species may also facilitate *S. aureus* hitchhiking into the growth medium through proximity to *C. albicans* cells emerging from the biofilm structure. It is well documented that *S. aureus* cell wall peptidoglycan is a potent inducer of hyphal formation [Xu], raising the possibility of peptidoglycan-driven enhancement of hyphal formation within cell clusters [Eichelberger and Cassat 2021)]. This may also promote additional opportunities for *S. aureus* attachment via Als3p and Als1p agglutinins [Peters *et al* (2012)]. These data suggest that study of yeast and hyphal-form biofilms by other methods, such as transcriptomic analysis, and further antimicrobial drug challenge over multiple time points during biofilm formation is required to confirm if morphotype specific clusters confer a spatial and/or temporal advantage in dual species biofilms. Our study tested the minimum concentration of vancomycin required to prevent planktonic growth, however, exposing yeast-form and hyphal form biofilms to increasing concentrations of vancomycin and at different time points would provide greater insight into the relationship of *C. albicans* cell morphotype, and highlight structurally-conferred tolerance to vancomycin.

These newly elucidated structural roles for *Candida* hyphae in the development of biofilms suggest that in addition to governing biofilm architecture, these structures may influence inherent antimicrobial resistance of a *C. albicans*/*S. aureus* dual species biofilm. If these data are reflected in the clinical setting, it is speculated that *C. albicans* cell morphotype plays a greater role in co-infection than simply a point of attachment and port of entry for systemic *S. aureus* infection [Ovchinnikova, Krom and Busscher (2012); Peters et al (2010)]. If *Candida* hyphae aid the nucleation of planktonic *S. aureus* cells *in vivo* to produce multiple large synergistic foci over a large surface area, as observed here, they may contribute to greater colonisation. Over the lifetime of the infection, this could have serious implications for treatment efficacy and patient outcomes.

## Acknowledgements

We would like to thank Professor Neil Gow and Professor Alistair Brown (both University of Exeter) for the gift of *C. albicans* strains and helpful discussions and to Dr Damir Sudar (Quantitative Imaging Systems LLC for the gift of a Q i Tissue license, and for discussions on analysis methods Thanks to Professor Carol Munro (University of Aberdeen), Dr Daniel Larcombe and Dr Liam Rooney (University of Strathclyde) for discussions on experimental design.

## Supplemental data

**Supplemental figure 1:**
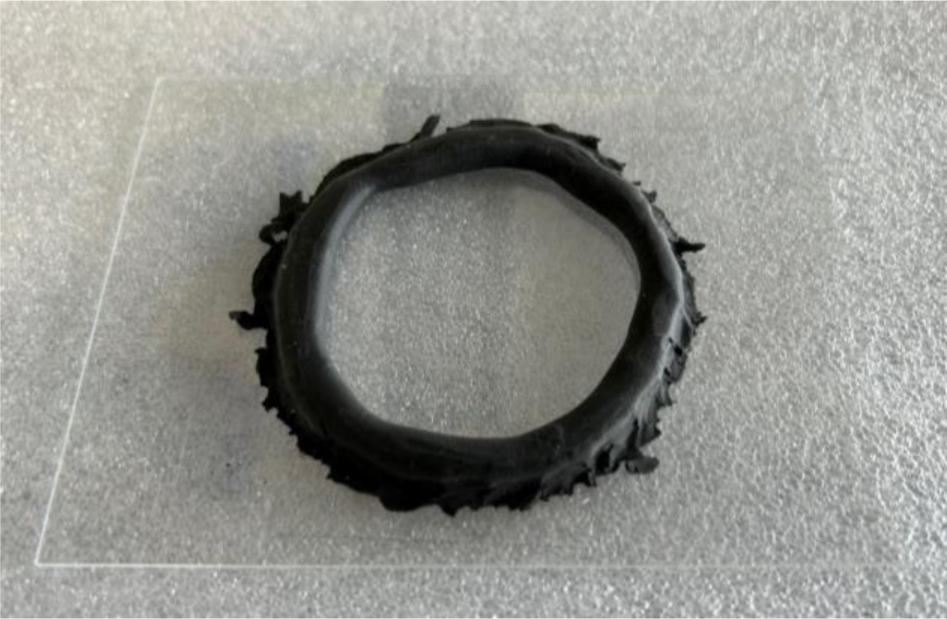
Image of Sugru dam manifold for time lapse mesoscopy. Hardened Sugru mouldable glue acts as a dam on a glass coverslip to retain agar without leakage, allowing agar to set into a thin layer for imaging purposes.

**Supplemental figure 2:**
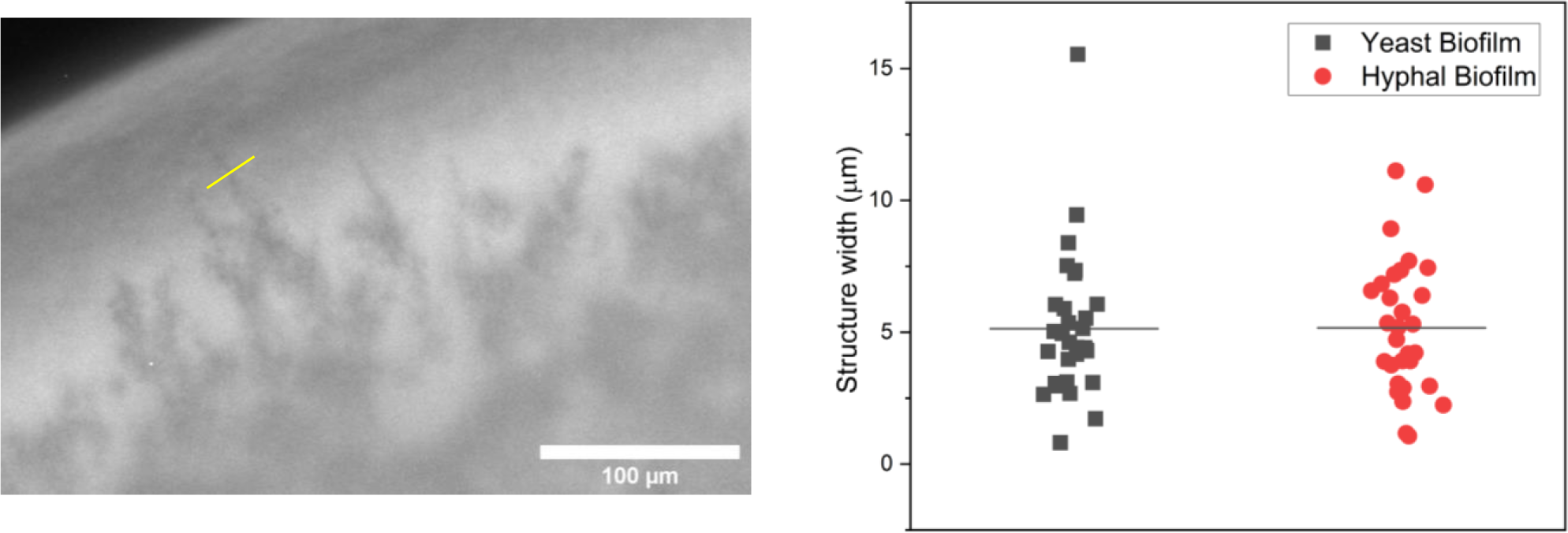
Measurement of *C.* albicans projections into *S. aureus* halo. **Panel A:** Illustration of measurement of *C. albicans* smallest projections using the Fiji line tool. Image is a maximum intensity z-projection from 9 hour time point. Scale bar 100µm. **Graph B:** Widths of *C. albicans* structures in both yeast-form and hyphal-form biofilm halos. Measurements taken from 3 independent experiments.

**Table 1.** Statistical analysis of quantification.

| Total diameter (μm) | Yeast-form Biofilm |  |  |  |  | Hyphal-form Biofilm |  |  |  |  |
| --- | --- | --- | --- | --- | --- | --- | --- | --- | --- | --- |
|  | Time point |  |  |  |  | Time point |  |  |  |  |
|  | t0 | t3 | t6 | t9 | t12 | t0 | t3 | t6 | t9 | t12 |
| Biofilm 1 | 3611.717 | 3670.111 | 3770.426 | 3909.015 | 4031.201 | 3722.796 | 3727.31 | 3844.39 | 3975.35 | 4129.101 |
| Biofilm 2 | 3334.269 | 3380.872 | 3486.628 | 3603.726 | 3770.23 | 3382.837 | 3385.088 | 3486.875 | 3619.118 | 3800.305 |
| Biofilm 3 | 3488.023 | 3537.235 | 3660.521 | 3810.563 | 3987.089 | 3705.838 | 3724.553 | 3872.435 | 4017.253 | 4170.505 |
| Average | 3478.003 | 3529.406 | 3639.192 | 3774.435 | 3929.507 | 3603.824 | 3612.317 | 3734.567 | 3870.574 | 4033.304 |
| Standard Deviation | 138.9951 | 144.7783 | 143.0962 | 155.8181 | 139.6899 | 191.5678 | 196.7909 | 214.9651 | 218.7726 | 202.842 |
| P value (T test) | 0.062886 | 0.133474 | 0.132004 | 0.11741 | 0.072096 |  |  |  |  |  |

| Core diameter (μm) | Yeast-form Biofilm |  |  |  |  | Hyphal-form Biofilm |  |  |  |  |
| --- | --- | --- | --- | --- | --- | --- | --- | --- | --- | --- |
|  | Time point |  |  |  |  | Time point |  |  |  |  |
|  | t0 | t3 | t6 | t9 | t12 | t0 | t3 | t6 | t9 | t12 |
| Biofilm 1 | 3174.842 | 3152.717 | 3017.968 | 3122.817 | 3165.587 | 3486.624 | 3442.392 | 3307.667 | 3221.460 | 3323.807 |
| Biofilm 2 | 2936.85 | 2898.258 | 2873.452 | 2932.953 | 2945.194 | 3198.962 | 3179.439 | 3071.307 | 3016.623 | 3094.862 |
| Biofilm 3 | 3088.439 | 3022.45 | 2922.811 | 3041.885 | 3109.019 | 3509.194 | 3485.348 | 3315.559 | 3279.723 | 3289.545 |
| Average | 3066.71 | 3024.475 | 2938.077 | 3032.552 | 3073.267 | 3398.26 | 3369.06 | 3231.511 | 3172.602 | 3236.071 |
| Standard Deviation | 120.4747 | 127.2416 | 73.45752 | 95.27548 | 114.4637 | 172.9657 | 165.6149 | 138.7968 | 138.1873 | 123.4849 |
| P value (T test) | 0.009695 | 0.014138 | 0.017444 | 0.05201 | 0.001588 |  |  |  |  |  |

| Halo width (μm) | Yeast-form Biofilm |  |  |  |  | Hyphal-form Biofilm |  |  |  |  |
| --- | --- | --- | --- | --- | --- | --- | --- | --- | --- | --- |
|  | Time point |  |  |  |  | Time point |  |  |  |  |
|  | t0 | t3 | t6 | t9 | t12 | t0 | t3 | t6 | t9 | t12 |
| Biofilm 1 | 188.16 | 249.48 | 334.607 | 390.255 | 433.626 | 117.367 | 142.167 | 250.782 | 347.263 | 405.062 |
| Biofilm 2 | 169.183 | 226.808 | 302.544 | 345.239 | 419.691 | 80.434 | 94.189 | 211.688 | 305.612 | 356.784 |
| Biofilm 3 | 181.767 | 239.388 | 326.449 | 386.335 | 436.138 | 102.989 | 113.190 | 267.144 | 357.576 | 407.678 |
| Average | 179.7033 | 238.5587 | 321.2 | 373.943 | 429.8183 | 100.2633 | 116.5153 | 243.2047 | 336.817 | 389.8413 |
| Standard Deviation | 9.655344 | 11.35873 | 16.66352 | 24.93554 | 8.860005 | 18.61675 | 24.16124 | 28.49393 | 27.51188 | 28.65836 |
| P value (T test) | 0.002124 | 0.001925 | 0.007351 | 0.00656 | 0.036659 |  |  |  |  |  |

